# ACETYLATED α-TUBULIN LYSINE 394 IS CRITICAL FOR AXON OUTGROWTH IN THE ADULT MUSHROOM BODIES OF *DROSOPHILA MELANOGASTER*

**DOI:** 10.1101/2025.10.10.681650

**Authors:** Chloe J. Welch, Liam O’Connor Mueller, Sophia P. Trujillo, Jill Wildonger

**Affiliations:** Biological Sciences Graduate Program, School of Biological Sciences, University of California, San Diego, 9500 Gilman Drive, La Jolla, CA 92093-0021, USA; School of Biological Sciences, Cell and Developmental Biology, University of California, San Diego, 9500 Gilman Drive, La Jolla, CA 92093-0021, USA; School of Biological Sciences, Molecular Biology, University of California, San Diego, 9500 Gilman Drive, La Jolla, CA 92093-0021, USA; School of Medicine, Pediatrics, University of California, San Diego, 9500 Gilman Drive, La Jolla, CA 92093-0021, USA

**Keywords:** *Drosophila melanogaster*, mushroom body, axon patterning, microtubule, Tau

## Abstract

Disruptions in the microtubule cytoskeleton play a role in various neurological diseases that afflict a large fraction of the population. Microtubule function is regulated by post-translational modifications like acetylation, and one consistently identified acetylation site in mammals and *Drosophila melanogaster* is α-tubulin lysine 394 (K394). Our previous research demonstrated that an acetylation-blocking point mutation—K394R—causes a decrease in microtubule stability in axon terminals at the developing neuromuscular junction. Here, we asked whether K394 acetylation regulates the development of additional neuronal structures. Using the central brain mushroom body as a model, we found that *K394R* results in β lobe overextension at the midline. The *K394R* phenotype manifests during metamorphosis and affects β lobe growth in a cell-autonomous manner. Our data suggest that the *K394R* phenotype may result from changes in Tau, a microtubule-associated protein enriched in the mushroom body and known to play a critical role in regulating neuronal microtubules. Knocking-out *tau* resulted in defects in midline crossing similar to *K394R*. However, when the loss of *tau* was combined with *K394R*, β lobe extension was normal—indicating that the loss of *tau* suppresses the *K394R* phenotype and vice versa. While overexpressing *tau* also resulted in a midline crossing phenotype, *K394R* in combination with elevated Tau resulted in a severely malformed mushroom body. Altogether, our work suggests that *K394R* interacts with *tau* to regulate axon outgrowth during mushroom body development and raises the potential of manipulating K394 acetylation to ameliorate neurological disease resulting from axonal growth defects and changes in Tau.

## INTRODUCTION

Developing neurons are highly dependent on the assembly, organization, and dynamic remodeling of microtubules, which are hollow, tube-like structures comprised of protofilaments of α- and β-tubulin heterodimers (Akhmanova and Steinmetz 2015; Kapitein and Hoogenraad 2015). Neuronal microtubule function is regulated by a variety of factors, including post-translational modifications (PTMs) and interactions with microtubule-associated proteins (MAPs), which influence microtubule formation, organization, and behavior (Janke and Bulinski 2011; Janke and Magiera 2020). PTMs affect microtubule stability in addition to the binding efficiency of MAPs and cargo trafficking mediated by molecular motors. One PTM involved in multiple aspects of neurodevelopment and function is acetylation (Kabir et al. 2022). Multiple lysine residues in tubulin—particularly in α-tubulin—are acetylated (L’Hernault and Rosenbaum 1985; Choudhary et al. 2009; Chu et al. 2011; Janke and Bulinski 2011; Janke and Magiera 2020). Although multiple tubulin acetylation sites have been identified, α-tubulin K40 is the only site that has been extensively characterized. Acetylation at K40 has historically been associated with stable microtubules in cells, but the PTM itself does not directly regulate dynamic microtubule properties; rather, previous studies have shown that this modification allows the microtubule cytoskeleton to endure mechanical stress, potentially explaining its affiliation with long-lived, stable microtubules across various cell types (Portran et al. 2017; Xu et al. 2017). However, whether and how microtubule function is influenced by acetylation of additional α-tubulin sites requires further investigation. Several studies have identified K394 as another conserved site of acetylation in multiple tissue types across many organisms, including tubulin from rodent brains and human cell culture lines (Choudhary et al. 2009; Weinert et al. 2011; Lundby et al. 2012; Liu et al. 2015a; Liu et al. 2015b; Weinert et al. 2018; Hansen et al. 2019). In contrast to K40, which is located within the lumen of the microtubule, K394 is located at the surface, at the α- and β-tubulin dimer interface, which makes it a promising candidate to regulate microtubule stability and interactions with proteins that influence microtubule function.

There are four α-tubulin isotypes encoded in the *Drosophila* genome, and the primary isotype, αTub84B, is ubiquitously expressed, is essential for survival, and shares over 95% identity with human α-tubulin (Raff 1984). In a previous study, we created a lysine (K) to arginine (R) point mutation in the αTub84B gene to block acetylation at the K394 site (K394R) (Saunders et al. 2022). Arginine and lysine have similar biochemical structure and charge, but arginine cannot be acetylated and therefore serves as a mimic for the loss of acetylation. In the same study, we showed that K394 acetylation can be detected in the *Drosophila* central nervous system and is critical for neuronal development and function by playing a role in axon terminal growth at the larval neuromuscular junction (NMJ). *K394R* results in a reduction of microtubule loops labeled by Futsch, a homolog of human Microtubule-Associated Protein 1B (MAP1B) that co-localizes with stable microtubules and influences synaptic growth (Hummel et al. 2000; Roos et al. 2000; Saunders et al. 2022). We also demonstrated that loss of histone deacetylase 6 (HDAC6) increases K394 acetylation levels, suggesting that this enzyme is the likely K394 deacetylase. Given the role of K394 acetylation in shaping axon terminal growth at the NMJ, it is likely that it is important for the development of additional structures in the fly nervous system.

During key periods of development, the growth of axons requires precise navigation toward and connection with their appropriate targets for proper wiring of the brain. Structural changes— such as synaptic changes or the misrouting of axons—are among the hallmark characteristics of various neurological diseases (Van Battum et al. 2015). Disruptions in axon guidance are common to neurodevelopmental disorders such as autism spectrum disorder (ASD) (McFadden and Minshew 2013; Van Battum et al. 2015). In individuals with ASD, abnormalities in axon size, thickness, and abundance have been documented in both the hippocampus and medial prefrontal cortex—structures of the brain that are critical for learning and memory formation in humans (Zikopoulos and Barbas 2010; Zikopoulos and Barbas 2013; Schubert et al. 2015). In the common fruit fly, *Drosophila melanogaster*, a structure called the mushroom body plays an analogous role in olfactory-based learning and memory and higher order sensory integration. The mushroom body is a powerful system to assess gene function in neurodevelopment— especially axon patterning (Lin 2023). The fly mushroom body is a bilaterally symmetric midbrain structure formed by a pair of neuropils containing approximately 2,000 neurons. The neurons of this structure, called Kenyon cells, are divided into subtypes based on their birth order and axonal projections. Kenyon cell axons extend medially and dorsally to form the α/β, α’/β’, and γ lobes. In wild-type flies, axons in the medially-projecting β lobes typically terminate near the midline of the brain but do not cross it. Disruptions in axon guidance are often characterized by the abnormal extension of β lobe axons across the midline. This β lobe midline crossing phenotype has been observed in *Drosophila* disease models including intellectual disability and Fragile X Syndrome, which is the most common, monogenic disorder associated with ASD (Michel et al. 2004; Kim et al. 2021; Stahl and Tomchik 2024; Genovese and Butler 2025).

In this study, we further explored the impacts of blocking α-tubulin K394 acetylation on axon morphogenesis by evaluating axon guidance during the development of the adult mushroom body. We found that the K394R mutants exhibit a β lobe midline crossing phenotype, which is first detectable during metamorphosis. Overexpressing *HDAC6* also causes overextension of the β lobes, indicating that the phenotype is a likely result of altering acetylation levels. Based on our previous findings that overexpressing Futsch suppresses the K394R phenotype at the larval NMJ (Saunders et al. 2022), we examined whether the mushroom body phenotype could be similarly suppressed by increasing the levels of Tau, a well-studied regulator of microtubule stability that is enriched in the mushroom body (Papanikolopoulou et al. 2019). Strikingly, we found that overexpressing *tau* in a *K394R* mutant background resulted in severe disruption in overall mushroom body morphology—a phenotype that was not observed in the *K394R* mutant alone. Knocking-out *tau* resulted in a β lobe midline crossing defect similar to the *K394R* mutant, and combining the *tau* knockout and *K394R* restored β lobe midline crossing levels to normal. These results point to a likely interaction between *tau* and K394 acetylation. Taken together, our data suggest that acetylation at K394 may affect Tau and its binding to microtubules and/or function to play an important role in axon morphogenesis in the developing mushroom body. Our results contribute to further characterizing conserved microtubule acetylation sites and support the use of *Drosophila* as a model for advancing our knowledge of how disruptions in microtubule regulators may be implicated in neurological disease.

## MATERIALS AND METHODS

### Fly stocks and husbandry

All fly crosses used for experiments were reared on molasses-yeast-agar food and maintained in a humidified incubator at 25°C on a constant 12:12 light/dark cycle. The following fly strains were obtained from the Bloomington Stock Center (Bloomington, IN, USA): *w^1118^*; *UAS-HDAC6* (#51181); *HDAC6^K/O^*(#51182); *OK107-GAL4* (#854); *UAS-mCD8::GFP* (#5137); *c305a-GAL4* (#30829); *201Y-GAL4* (#4440); *UAS-mCD8::GFP, c739-GAL4* (#64305); *tau^K/O^* (#64782). The *UAS-tau::HA* line was a generous gift from Dr. Sebastian Rumpf (University of Münster, North Rhine-Westphalia, Germany) (Herzmann et al. 2018). The following fly strains were described previously: *αTub84B^K394R^* and *UAS-αTub84B* (Saunders et al. 2022).

### Immunohistochemistry and confocal microscopy

Fasciculating axons in the mushroom body were visualized using an adaptation of previously established methods (Michel et al. 2004; Nguyen et al. 2021). Flies aged 0-2 days were dissected in 1% normal goat serum in 1X phosphate-buffered saline (1% NGS-PBS), then fixed in 4% paraformaldehyde (PFA) for 25 minutes at room temperature (RT). One quick wash was performed, then brains were washed three times for 15 minutes each in 0.1% PBS in 10% Triton X-100 (0.1% PBST) at RT. Brains were then blocked in 1% bovine serum albumin (BSA) in PBST for 1 hour at RT, and incubated in primary antibody overnight at 4°C (*α*/β neurons: mouse anti-Fasciclin II [Developmental Studies Hybridoma Bank, Iowa City, IA] at 1:20 in 0.1% PBST; α’/β’/γ neurons: mouse anti-Trio [Developmental Studies Hybridoma Bank, Iowa City, IA] at 1:20 in 0.1% PBST). Brains were washed three times for 20 minutes each in 0.1% PBST at RT, then incubated in secondary antibody overnight at 4°C in the dark (goat anti-mouse AlexaFluor® 488 [Jackson ImmunoResearch] at 1:1000 in 0.1% PBST). Brains were washed six times for 20 minutes each in 0.1% PBST at RT and mounted in Prolong® Gold antifade mountant (Invitrogen). Z-stack images of all mushroom bodies were captured using a Leica Stellaris LSM confocal microscope. For late third instar larvae, a mix of male and female brains were analyzed. For 2-3 days after pupal formation (APF) stages and for 0-2 day old adult stages, only male brains were analyzed.

### Quantification and statistical analyses

All image analyses were performed using FIJI/ImageJ, and all statistical analyses were performed using Prism 10 (GraphPad Software, San Diego, CA) and R (R Core Team 2025). Images were blinded prior to analysis. Whole percentages of phenotype expressed were calculated for each category of crossing severity (number of brains per score/total sample size for given genotype). The frequency and severity of mushroom body β lobe midline crossing was scored manually as either no crossing, partial crossing, or complete crossing (adapted from Michel et al. 2004). In order to demonstrate that partial and complete crossing are distinct groups in our analysis, representative images from each category were selected and axon fiber thickness was quantified (microns) in Fiji/ImageJ and a Student’s t-test with Welch’s Correction was run (Table S1). To test the null hypothesis that distinct genotypes had similar probabilities of expressing either the “partial crossing” or “complete crossing” phenotype, generalized linear models (GLM) with a binomial error structure were built in R. Model fit, a common concern in GLMs, was deemed to not be an issue as residual deviance was always close to the degrees of freedom in these models (McCullagh and Nelder 1989). Bars on graphs represent probability ± 95% confidence intervals (CIs) extracted from the GLMs. As these models cannot converge when no extreme phenotypes are observed, a Fisher’s exact test was performed in those cases. For all tests, an alpha value of *p ≤* 0.05 was used to determine statistical significance.

## RESULTS

### Disruption of α-tubulin K394 acetylation causes defects in mushroom body β lobe extension at the midline

The *Drosophila* mushroom body is comprised of bilaterally symmetric, paired neuropils divided into specific anatomical domains (Lin 2023). The Kenyon cell bodies are clustered in the dorsal posterior cortex of the central brain. The Kenyon cell dendrites form a calyx beneath the cell bodies, and the Kenyon cell axons extend toward the surface of the brain where they form a peduncle made of five terminal lobes, referred to as the α, α’, β, β’, and γ lobes. The mushroom body neurons are generated sequentially during development, and axons from the earliest born neurons first establish the medial γ lobe. The neurons born next project axons that bifurcate to form the dorsal α’ lobe and medial β’ lobe, and the axons of the neurons born last also bifurcate, forming the dorsal α lobe and medial β lobe (Fig. 1a). The medial β, β’, and γ lobes terminate near the midline of the adult brain.

**Figure 1.**
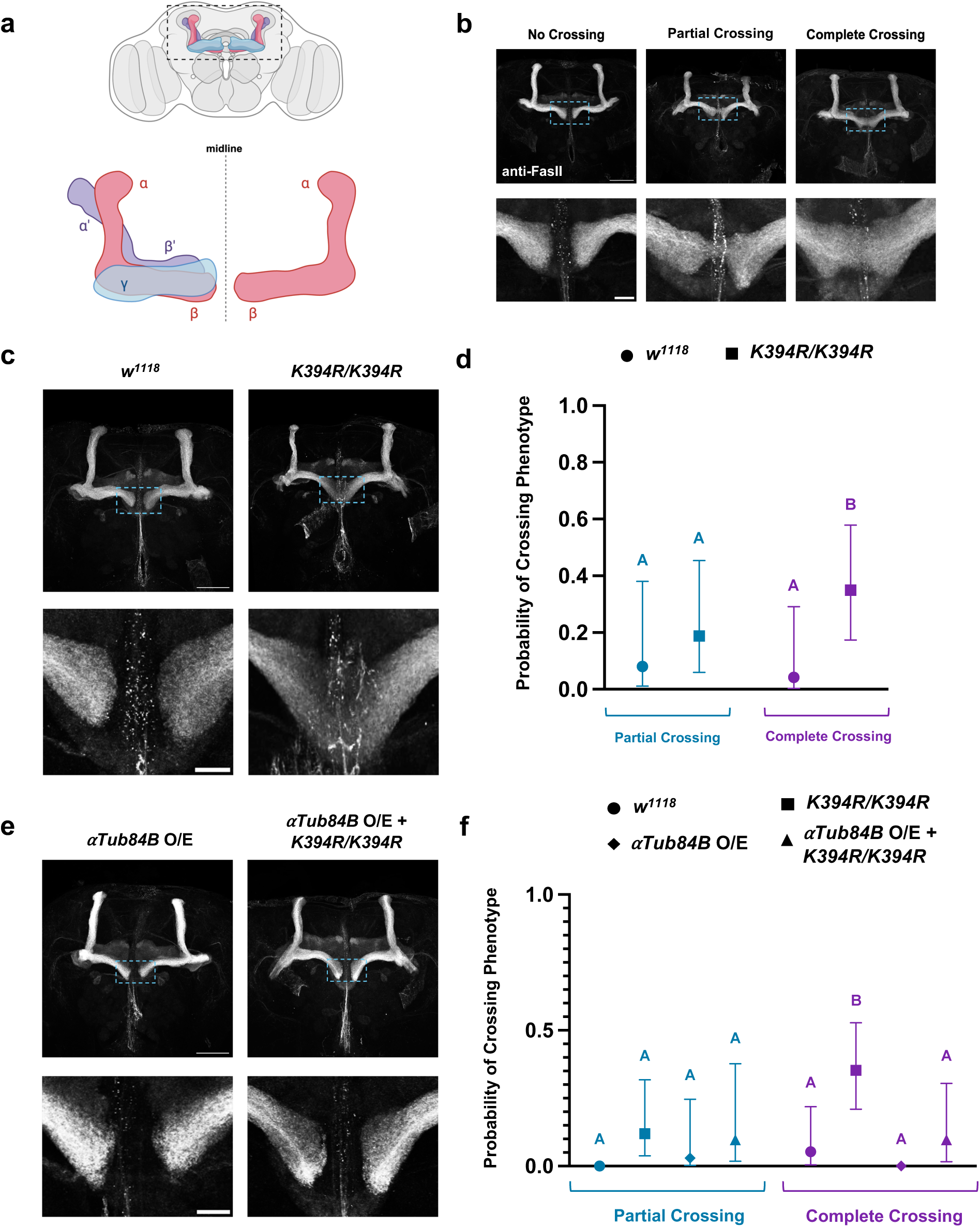
Disruption of K394 acetylation causes defects in mushroom body β-lobe extension at the midline. **(a)** Cartoon schematic of adult *Drosophila* brain showing different lobes of the mushroom body. Blue = γ lobes; Pink = α/β lobes; Purple = α’/β’ lobes. **(b)** Representative images of mushroom bodies stained with anti-Fasciclin II (anti-FasII)—which stains the α/β lobes—highlighting three categories of midline crossing ranging in severity. **(c)** Representative images of *w^1118^* control or *K394R/K394R* mutant mushroom bodies. **(d)** Quantification of crossing frequencies as probabilities of either (1) Partial Crossing or (2) Complete Crossing phenotypes being different from *w^1118^* control mushroom bodies. The frequency of complete crossing occurring was higher in *K394R* homozygous mutant mushroom bodies. Sample sizes: *w^1118^* = 26; *K394R/K394R* = 23. **(e)** Representative mushroom body images of animals overexpressing wild-type *αTub84B (αTub84B O/E*) or overexpressing wild-type *αTub84B* in the *K394R/K394R* mutant *(αTub84B O/E* + *K394R/K394R*). **(f)** Quantification of crossing frequencies as probabilities of either (1) Partial Crossing or (2) Complete Crossing phenotypes being different from *w^1118^*control mushroom bodies. Overexpression of wild-type *αTub84B* in *K394R* homozygous background reverts midline crossing phenotype. Sample sizes: *w^1118^* = 38*; K394R/K394R* = 37; *αTub84B O/E* = 34; *αTub84B O/E* + *K394R/K394R* = 34. Values at “0” indicate phenotype was not observed. Bars on graphs represent 95% confidence intervals (upper/lower ranges) for probabilities. Statistical significance indicated by different letters where “Partial Crossing” and “Complete Crossing” are treated independently from one another. Blue dashed boxes represent zoomed-in views of mushroom body midline. Scale bars: 50 μm, 10 μm.

To determine whether acetylation of α-tubulin K394 affects axon patterning in the brain, we first examined how the *K394R* mutation impacts mushroom body development—particularly β lobe axon extension at the midline of the brain. β lobe axon extension phenotypes were divided into three categories: (1) “no crossing,” in which β lobe axons terminate before the midline; (2) “partial crossing,” in which a few axons extend across the midline; or (3) “complete crossing,” in which the β lobe axons project across the midline, forming a thick bundle of axons between the β lobes in each hemisphere (Fig. 1b). We utilized an antibody against the cell adhesion molecule Fasciclin II (FasII), as it strongly labels the α and β lobes of the adult mushroom body, and only weakly labels the γ lobes (Crittenden et al. 1998). In *w^1118^* control brains, β lobe axons terminated near the midline and overextension was rarely observed. In contrast, in the *K394R* homozygous mutant brains, partial crossing was observed in nearly 20% of mushroom bodies and complete crossing was observed in 35% of mushroom bodies (Fig. 1, c and d). This suggests that K394 acetylation plays a role in axon patterning in the mushroom body—building on our previous work demonstrating its role in shaping axon terminal growth at the NMJ.

We next tested whether the midline crossing defect observed in the *K394R* homozygous mutants was due to cell-autonomous or non-autonomous disruption of α-tubulin and microtubules. To do this, we expressed transgenic wild-type *α-Tub84B* (*UAS-αTub84B*) specifically in the mushroom body using the mushroom body-specific driver *OK107-GAL4* in the *K394R* mutant background (*αTub84B O/E + K394R/K394R*). In the mushroom bodies of *αTub84B O/E + K394R/K394R* flies, we found that β lobe axon extension was restored to normal (Fig. 1, e and f), with the frequency of complete crossing occurring in less than 10% of mushroom bodies—similar to *w^1118^* controls and in contrast to the ∼35% observed in the *K394R* homozygous mutants. These results suggest that the midline crossing phenotype is a result of the *K394R* mutation acting cell-autonomously and that K394 acetylation is critical for proper axon outgrowth in the adult mushroom body.

### Overexpression of *HDAC6* results in a midline crossing phenotype

To further test whether the *K394R* phenotypes result from the loss of K394 acetylation, we over-expressed HDAC6, which our previous work indicates is the K394 deacetylase (the enzyme that acetylates K394 is not known) (Saunders et al. 2022). We over-expressed *HDAC6* in the mushroom body (*UAS-HDAC6; OK107-GAL4*) and found that *HDAC6* overexpression (*HDAC6 O/E*) resulted in ∼15-20% of mushroom bodies displaying either partial crossing or complete crossing (Fig. 2, a and b). The overexpression of *HDAC6* in combination with *K394R* did not result in a more severe phenotype than either alone, consistent with the idea that *HDAC6* and *K394R* are functioning in the same pathway to regulate β lobe extension (Fig. 2, a and b).

**Figure 2.**
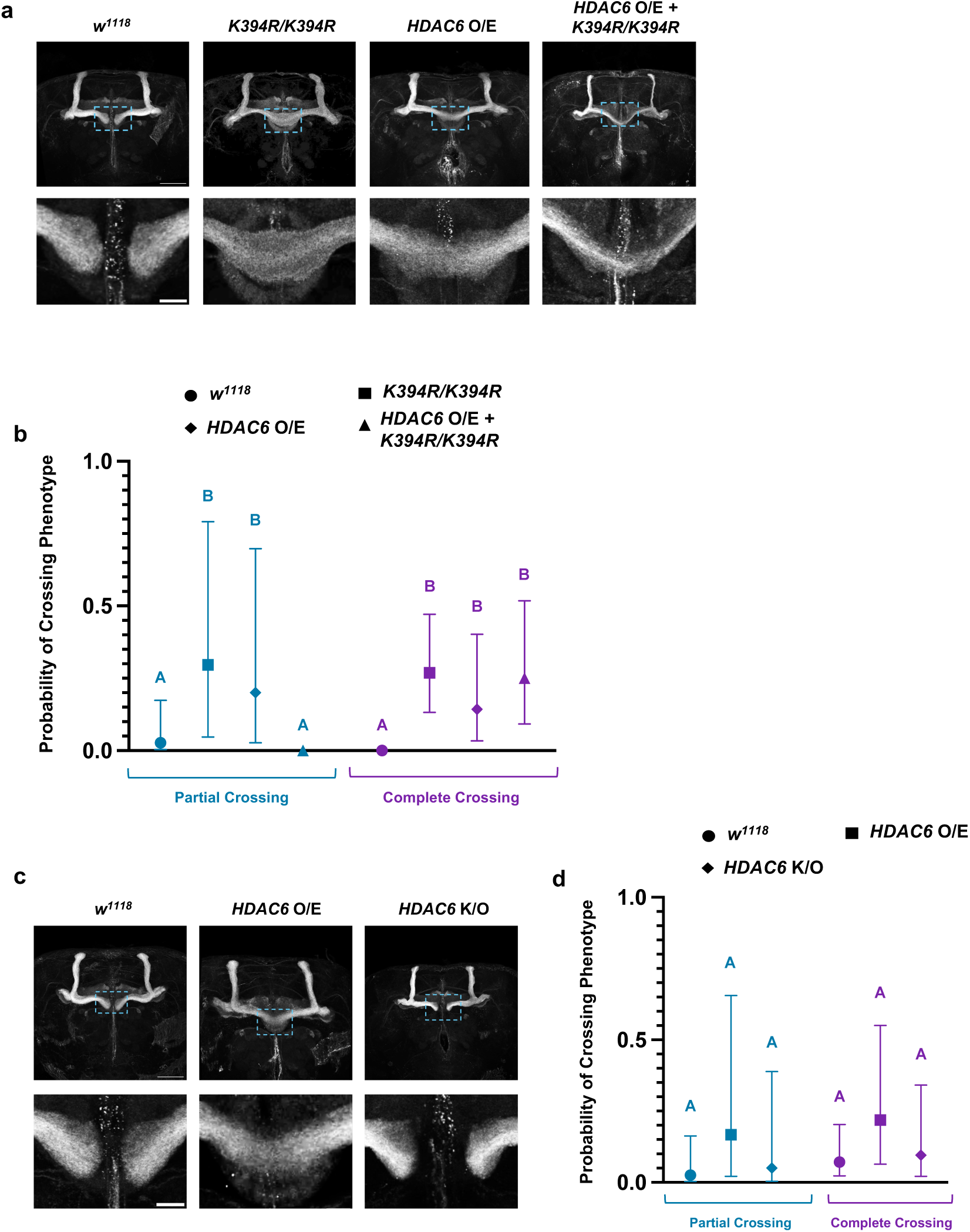
Overexpression of *HDAC6* results in a midline crossing phenotype. (a) Representative images of *w^1118^* control mushroom bodies, *K394R/K394R* mutant mushroom bodies, *HDAC6 O/E* mushroom bodies, and mushroom bodies of animals overexpressing wild-type *HDAC6* in the *K394R/K394R* mutant (*HDAC6 O/E* + *K394R/K394R*). **(b)** Quantification of crossing frequencies as probabilities of either (1) Partial Crossing or (2) Complete Crossing phenotypes being different from *w^1118^* control mushroom bodies. The frequency of Partial Crossing occurring was higher in the *K394R* homozygous mutant and *HDAC6 O/E* brains, and the frequency of Complete Crossing occurring was higher in the *K394R* homozygous mutant, *HDAC6 O/E*, and *HDAC6 O/E* + *K394R* brains. Sample sizes: *w^1118^*= 37; *K394R/K394R* = 34; *HDAC6 O/E* = 34; *HDAC6 O/E* + *K394R/K394R* = 36. **(c)** Representative images of *w^1118^* control mushroom bodies, mushroom bodies of animals overexpressing wild-type *HDAC6* (*HDAC6 O/E*), and mushroom bodies of animals homozygous for an *HDAC6* knockout allele (*HDAC6 K/O*). **(d)** Quantification of crossing frequencies as probabilities of either (1) Partial Crossing or (2) Complete Crossing phenotypes being different from *w^1118^* control mushroom bodies. Sample sizes: *w^1118^* = 43; *HDAC6 O/E* = 37; *HDAC6 K/O* = 44. Values at “0” indicate phenotype was not observed. Bars on graphs represent 95% confidence intervals (upper/lower ranges) for probabilities. Statistical significance indicated by different letters where “Partial Crossing” and “Complete Crossing” are treated independently from one another. Blue dashed boxes represent zoomed-in views of mushroom body midline. Scale bars: 50 μm, 10 μm.

Next, we asked whether the loss of HDAC6, which is predicted to result in an increase in K394 acetylation, has any effect on mushroom body development. In contrast to the overexpression of *HDAC6*, β lobe extension was normal in flies homozygous for a knockout allele of *HDAC6* (*HDAC6 K/O*) (Fig. 2, c and d). In this experiment, the overexpression of *HDAC6* increased the frequency of midline crossing (15-20%), but the result was not significant, indicating that the effect of *HDAC6* overexpression may be weak and less robust than the effect of the *K394R* mutation (Fig. 2, c and d). It is possible that elevating the levels of HDAC6 is not as efficient as mutating K394 in blocking acetylation at this site. Altogether, these experiments with *HDAC6* are consistent with the model that loss of K394 acetylation disrupts the extension of the β lobes.

### β lobe midline crossing phenotype in the *K394R* mutant manifests during metamorphosis

Our data shows that *K394R* causes defects in β lobe axon outgrowth. This led us to next determine when the β lobe overextension phenotype manifests during mushroom body development. The mushroom body β lobe neurons are born ∼one day after pupal formation. Marked by FasII, β lobe formation was clear at 2-3 days after pupal formation (APF) (Fig. 3, a and b; Table 1). We observed complete midline crossing at this stage in the *K394R* homozygous mutants, indicating that these axon pathfinding defects are evident during metamorphosis (Fig. 3b; Table 1).

**Figure 3.**
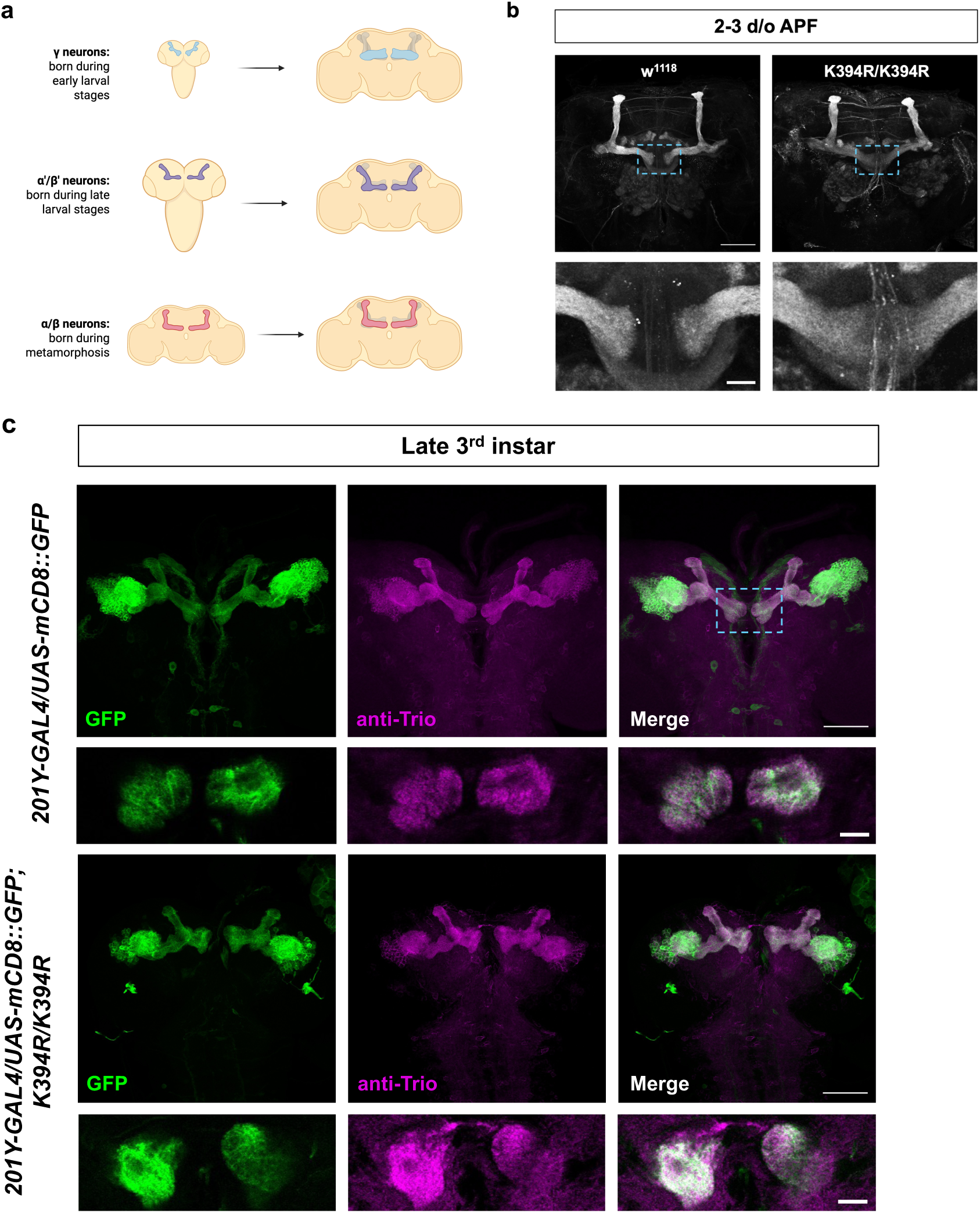
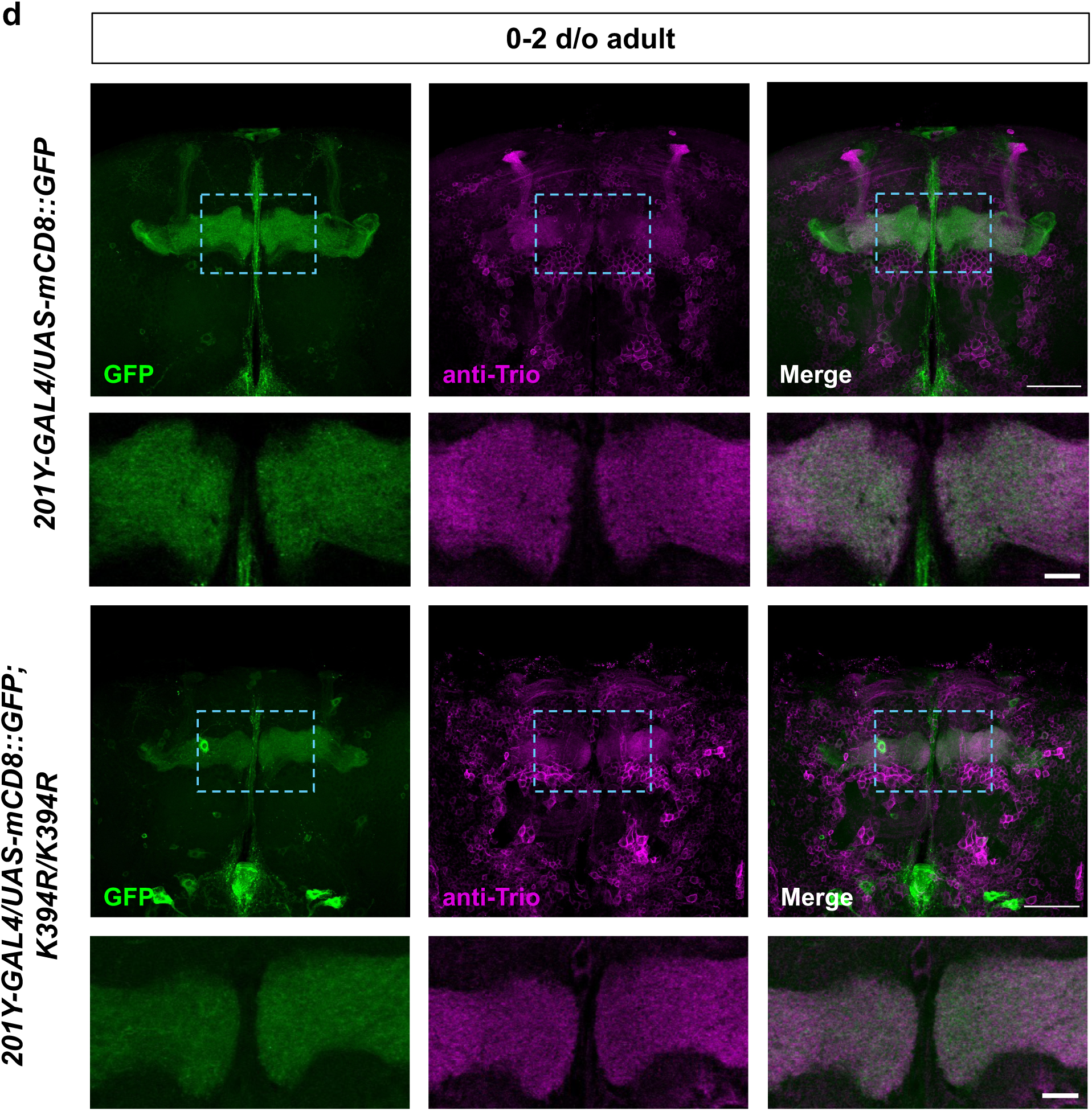
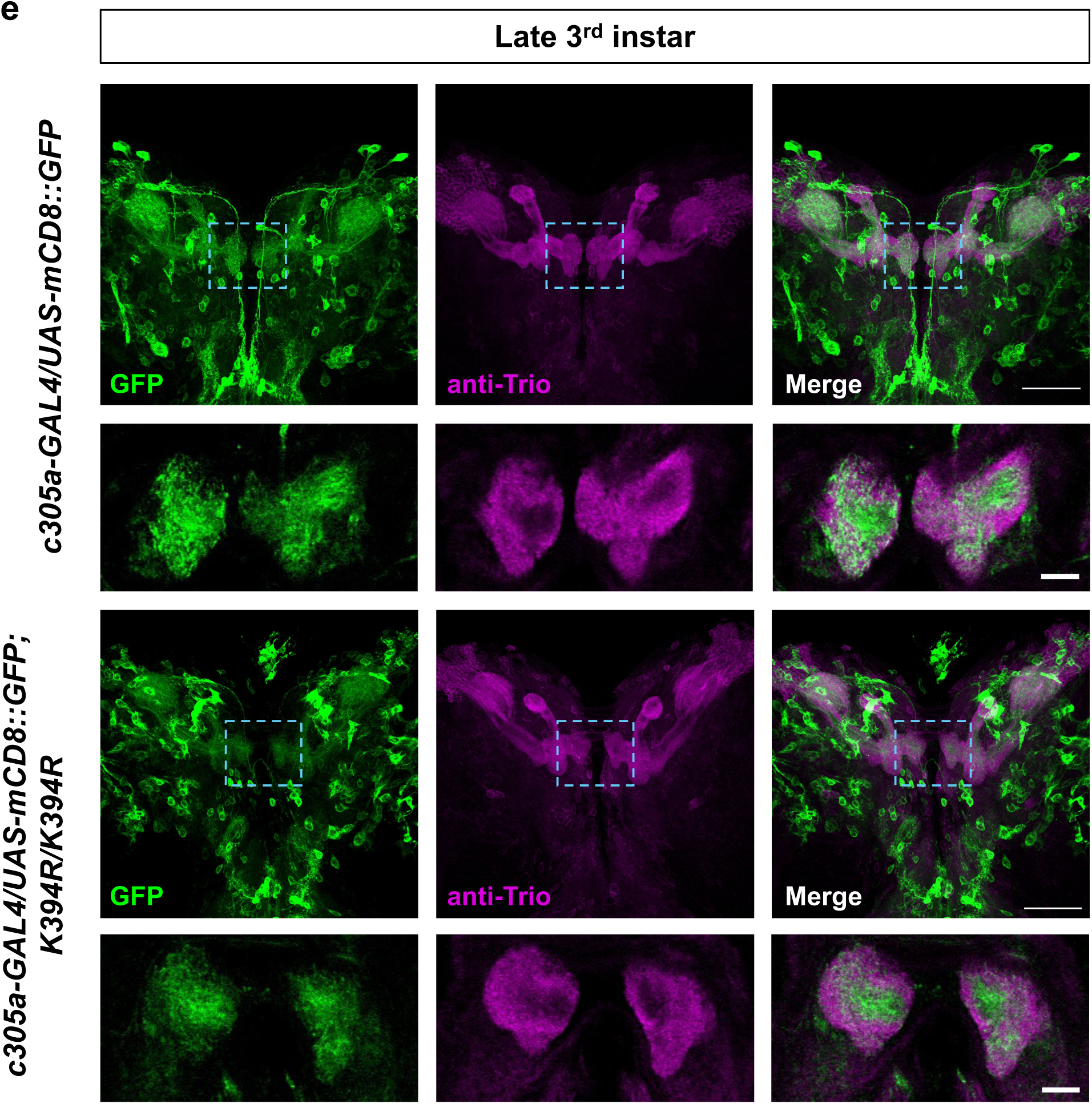
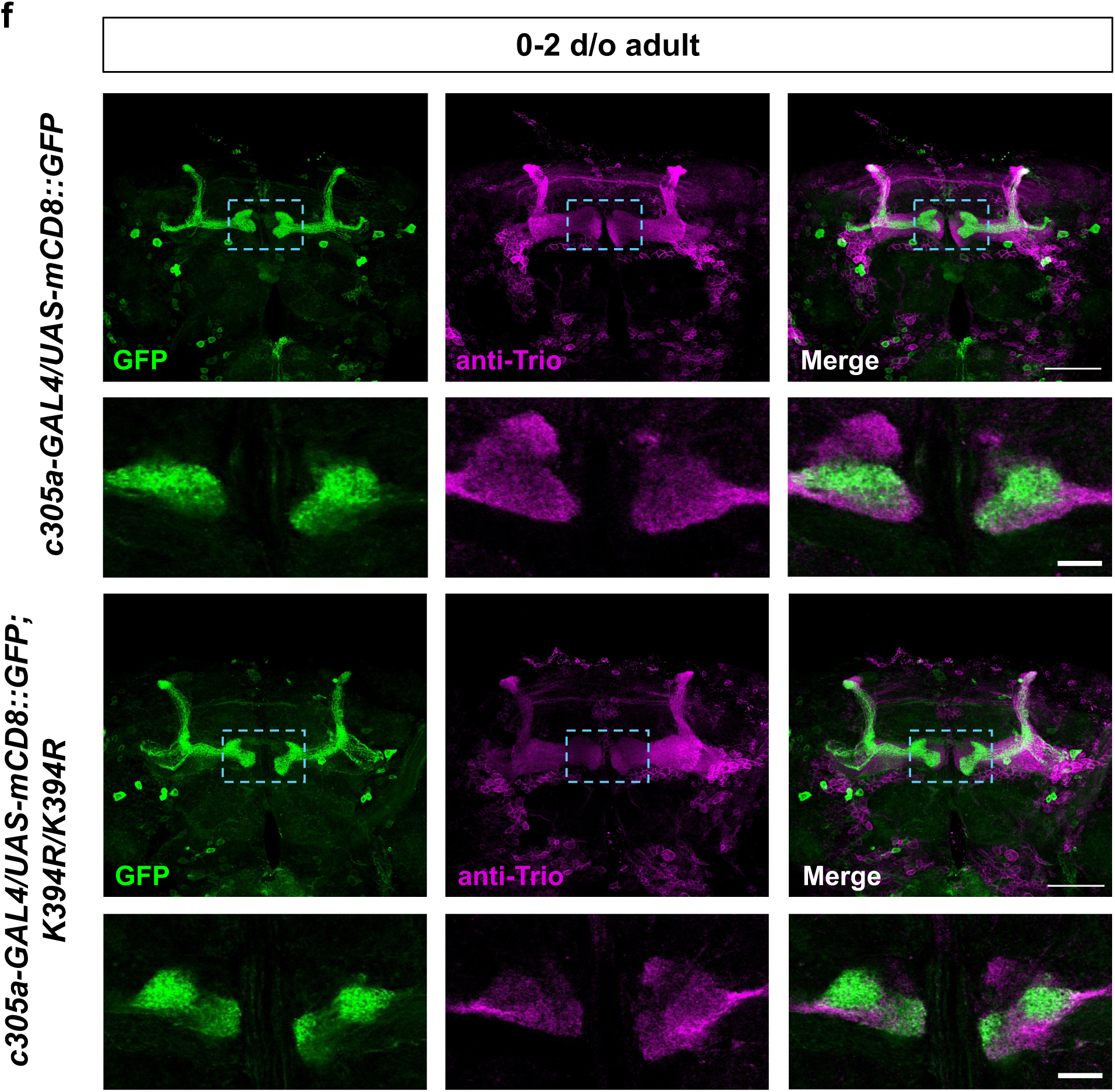
β-lobe midline crossing phenotype in the *K394R* mutant manifests during metamorphosis. **(a)** Schematic illustrating formation of mushroom body lobes during *Drosophila* development. γ neurons are born first during the early larval stages, followed by α’/β’ neurons during the late larval stages, and α/β neurons born during metamorphosis. **(b)** Representative images of *w^1118^* control and *K394R/K394R* mutant mushroom bodies at 2-3 days after pupal formation (APF). Mushroom bodies were labeled with anti-FasII antibody to visualize α/β lobes. Midline crossing can be detected during this stage in *K394R/K394R* mutants. **(c-d)** Representative images of mushroom bodies of animals expressing *UAS-mCD8::GFP* under control of the γ-neuron driver, *201Y-GAL4*, by itself and in the *K394R/K394R* mutant at the late 3^rd^ instar larval stage (c) and at the adult stage (d). **(e-f)** Representative images of mushroom bodies of animals expressing *UAS-mCD8::GFP* under control of the α’/β’-neuron driver, *c305a-GAL4*, by itself and in the *K394R/K394R* mutant at the late 3^rd^ instar larval stage (e) and at the adult stage (f). Mushroom bodies were co-labeled with anti-Trio antibody, which stains the γ and α’/β’ lobes of the mushroom body. Sample sizes: *w^1118^* = 33; *K394R/K394R* = 21 (b); *201Y-GAL4/UAS-mCD8::GFP* = 28 (c) and 34 (d); *201Y-GAL4/UAS-mCD8::GFP*; *K394R/K394R* = 21 (c) and 26 (d); *c305a-GAL4/UAS-mCD8::GFP* = 41 (e) and 32 (f); *c305a-GAL4/UAS-mCD8::GFP*; *K394R/K394R* = 24 (e) and 34 (f). Blue dashed boxes represent zoomed-in views of mushroom body midline. Zoomed-in images: single slice. Scale bars: 50 μm, 10 μm.

**Table 1.** Midline crossing phenotype is detectable at 2-3 days APF in *K394R* mutants. No midline crossing was observed between γ lobes or α’/β’ lobes, indicating that lobe overextension is specific to the β lobes and manifests during metamorphosis. For each category of crossing, total numbers of brains are indicated.

| GENOTYPE | DEVELOPMENTAL STAGE | NO CROSSING | PARTIAL CROSSING | COMPLETE CROSSING |
| --- | --- | --- | --- | --- |
| *201Y-GAL4/*<br>*UAS-mCD8::GFP* | Late 3<sup>rd</sup> instar | 28 | 0 | 0 |
| *201Y-GAL4/*<br>*UAS-mCD8::GFP;*<br>*K394R/K394R* | Late 3<sup>rd</sup> instar | 21 | 0 | 0 |
| *201Y-GAL4/*<br>*UAS-mCD8::GFP* | 0-2 d/o adult | 34 | 0 | 0 |
| *201Y-GAL4/*<br>*UAS-mCD8::GFP;*<br>*K394R/K394R* | 0-2 d/o adult | 26 | 0 | 0 |
| *c305a-GAL4/*<br>*UAS-mCD8::GFP* | Late 3<sup>rd</sup> instar | 41 | 0 | 0 |
| *c305a-GAL4/*<br>*UAS-mCD8::GFP;*<br>*K394R/K394R* | Late 3<sup>rd</sup> instar | 24 | 0 | 0 |
| *c305a-GAL4/*<br>*UAS-mCD8::GFP* | 0-2 d/o adult | 32 | 0 | 0 |
| *c305a-GAL4/*<br>*UAS-mCD8::GFP;*<br>*K394R/K394R* | 0-2 d/o adult | 34 | 0 | 0 |
| *w<sup>1118</sup>* | 2-3 days APF | 33 | 0 | 0 |
| *K394R/K394R* | 2-3 days APF | 17 | 0 | 4 |

The α/β neurons are the last mushroom body neurons to be born, raising the possibility that the misrouting of axons from earlier born neurons might underlie the β lobe midline crossing phenotype. The first mushroom body axons to extend are the γ lobe axons, followed by the axons of the α’/β’ lobes (Fig. 3a) (Lee et al. 1999). To establish whether the γ or α/β lobes are affected in the *K394R* mutant, we performed a set of temporal development experiments and analyzed each lobe subset at two different time points. We focused our analysis on the γ, β’, and β lobes as the midline crossing phenotype is specific to the medially-projecting lobes. To assess γ lobe development, we fluorescently labeled the neurons with γ lobe-specific *201Y-GAL4 UAS-mCD8::GFP* and used an antibody against Trio, which marks the γ and α’/β’ lobes (Fig. 3, c and d). To evaluate β’ lobe development, we used anti-Trio and *c305a-GAL4 UAS-mCD8::GFP*, which specifically illuminates the α’/β’ lobes (Fig. 3, e and f; Table 1). We scored the mushroom bodies in control and *K394R* mutant brains from late third instar larvae and 0-2 day old (d/o) adults, and no overextension of the γ lobes or β’ lobes was observed at either the late third instar or adult stage in *K394R* mutant mushroom bodies (Fig. 3, c-f; Table 1). Altogether, these data suggest that the axon overextension phenotype in the *K394R* mutant manifests as the β lobes develop during pupal stages and that the development of the γ and α’/β’ lobes are not overtly affected. Thus, the *K394R* mutant phenotype is specific to the β lobes and does not arise from obvious changes in the axonal projections of earlier born neurons.

### *K394R* in combination with elevated *tau* expression severely disrupts α and β lobe development

Our previous studies examining the NMJ in *K394R* mutant larvae revealed that the mutation disrupts microtubule stability, making microtubules more sensitive to destabilization by nocodazole (Saunders et al. 2022). Increasing microtubule stability, via the microtubule stabilizing agent taxol, or by elevating the levels of the MAP Futsch, suppressed the *K394R* NMJ phenotypes. Given these data, we asked whether a microtubule regulator that is enriched in the mushroom body, Tau, might similarly revert the *K394R* phenotype in this structure (Papanikolopoulou et al. 2019). We found that the overexpression of *tau* in the mushroom body (*OK107-GAL4 UAS-tau*, or *tau O/E*) resulted in overextension of the β lobes, with ∼27% of mushroom bodies showing complete crossing (Fig. 4, a and b). Tau is typically thought to positively regulate microtubule stability, leading us to expect that elevating tau levels would suppress the *K394R* mushroom body phenotype, analogous to our finding that elevating Futsch suppresses the *K394R* larval NMJ phenotypes (Saunders et al. 2022). However, our results showed that overexpressing *tau* in the *K394R* mutant background (*tau O/E + K394R/K394R*) severely disrupted the overall development of the α and β lobes in ∼90% of the brains (Fig. 4, a and c). The α and β lobes were either abnormal in appearance, with dramatically truncated lobes, or completely absent (Fig. 4a). Abnormal α and β lobes were not observed in control or *K394R* mutant brains, although ∼20% of *tau O/E* brains had abnormal α and β lobes (Fig. 4, a and b). Our results suggest that the *K394R* mutation significantly enhances the effect of elevated Tau on α and β lobe development.

**Figure 4.**
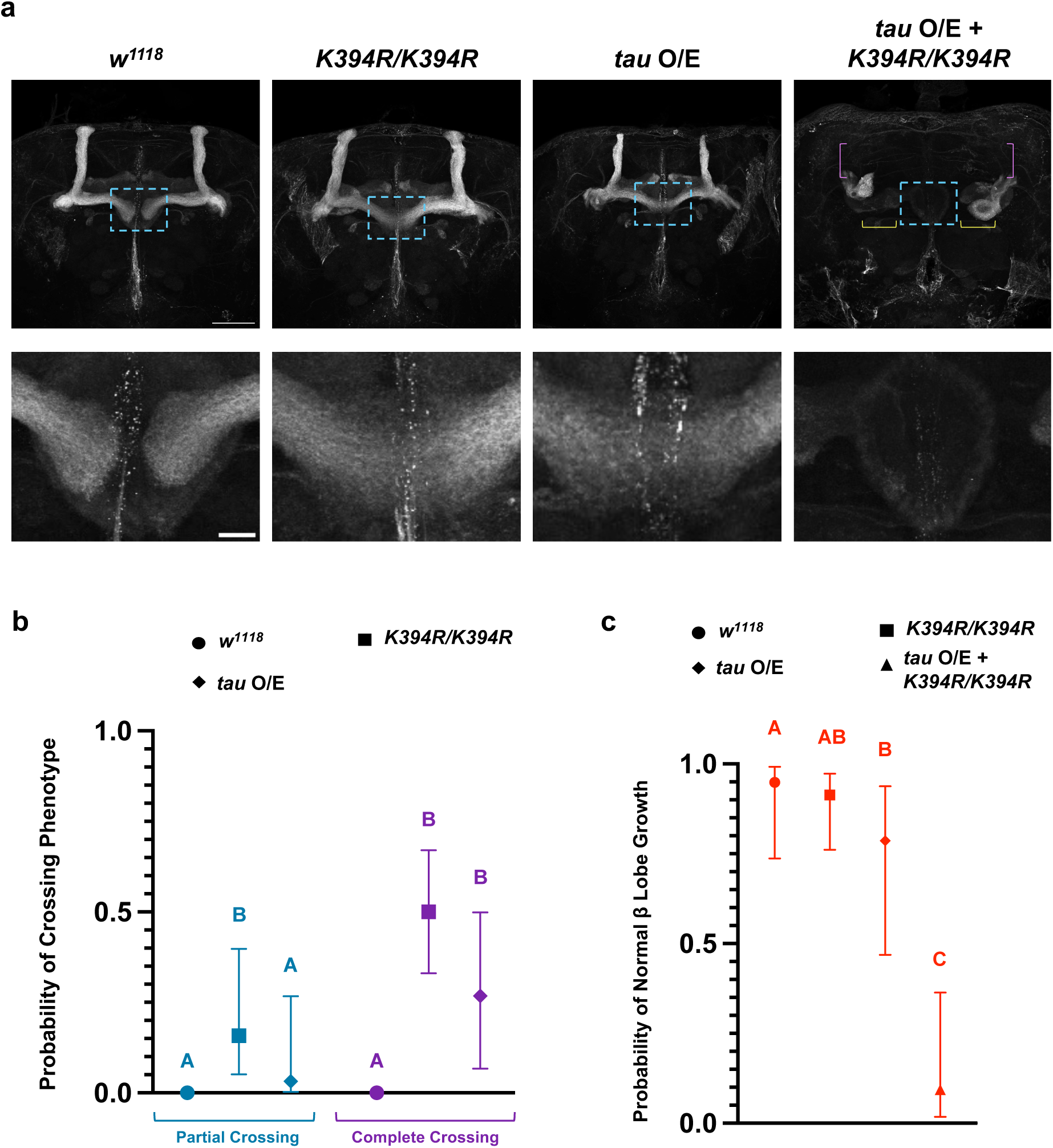
*K394R* enhances β-lobe midline crossing phenotype caused by *tau* overexpression. **(a)** Representative images of *w^1118^* control mushroom bodies, *K394R/K394R* mutant mushroom bodies, mushroom bodies of animals overexpressing wild-type *Drosophila tau* (*tau O/E*), and mushroom bodies of animals overexpressing *tau* in the *K394R/K394R* mutant (*tau O/E* + *K394R/K394R*). *Tau O/E* + *K394R/K394R* caused severe defects in overall mushroom body morphology. **(b)** Quantification of crossing frequencies as probabilities of either (1) Partial Crossing or (2) Complete Crossing phenotypes being different from *w^1118^*control mushroom bodies. The frequency of Complete Crossing occurring was higher in *tau O/E* mushroom bodies compared to *w^1118^* controls. **(c)** Quantification of mushroom bodies showing normal β lobe growth (no lobes missing or abnormal in size) as probabilities compared across all genotypes. The frequency of mushroom bodies exhibiting normal β lobe growth was lower in *tau O/E* brains compared to *w^1118^*but not *K394R/K394R*, and lower in *tau O/E* + *K394R/K394R* brains compared to *w^1118^* and *K394R/K394R*. Sample sizes: *w^1118^* = 39; *K394R/K394R* = 35; *tau O/E* = 42; *tau O/E* + *K394R/K394R* = 32. Purple brackets indicate severely truncated α lobes. Yellow brackets indicate severely truncated/missing (abnormal) β lobes. For (b): Bars on graphs represent 95% confidence intervals (upper/lower ranges) for probabilities. Statistical significance indicated by different letters where “Partial Crossing” and “Complete Crossing” are treated independently from one another. For (c): Bars on graphs represent 95% confidence intervals (upper/lower ranges) for probabilities. Statistical significance indicated by different letters. Purple brackets indicate severely truncated α lobes. Yellow brackets indicate severely truncated/missing (abnormal) β lobes. Blue dashed boxes represent zoomed-in views of mushroom body midline. Scale bars: 50 μm, 10 μm.

### *K394R* suppresses the effects of knocking out *tau* on β lobe midline overextension

Our striking result that *K394R* enhances the deleterious effects of evaluated Tau on α and β lobe development suggests that *K394R* and *tau* interact. To evaluate a potential genetic interaction between *K394R* and *tau*, we examined the mushroom bodies of flies homozygous for a *tau* knockout in combination with *K394R* (*tau^K/O^, K394R/tau^K/O^, K394R*) and compared them to the mushroom bodies in homozygous *K394R* and *tau^K/O^*single mutants. Similar to the *K394R* mutants, the *tau^K/O^*mushroom bodies displayed a significant β lobe midline crossing phenotype, with ∼75% of the mushroom bodies showing complete crossing (Fig. 5, a and b). In the double-mutant *tau^K/O^, K394R* brains, β lobe midline crossing frequencies were restored to normal (Fig. 5, a and b). Thus, eliminating *tau* suppresses the effects of the *K394R* mutation on mushroom body development. These data are consistent with the idea that there is an interaction between *K394R* and *tau* that is critical for proper axon outgrowth to generate a mushroom body in the adult fly central nervous system.

**Figure 5.**
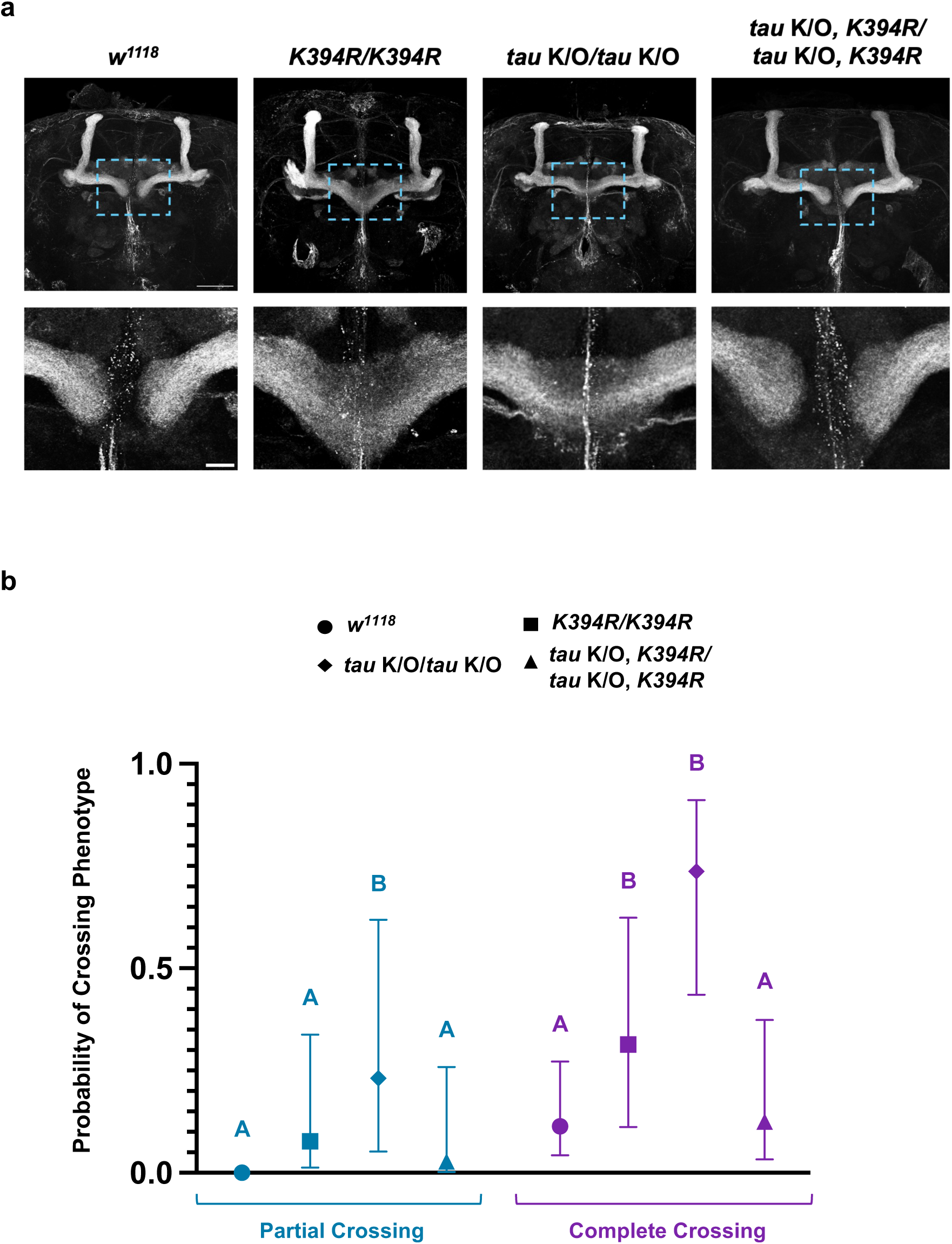
*K394R* suppresses the effects of knocking-out *tau* on β-lobe midline overextension. **(a)** Representative images of *w^1118^* control mushroom bodies, *K394R/K394R* mutant mushroom bodies, *tau^K/O^*/*tau^K/O^* mushroom bodies, and mushroom bodies of homozygous animals with *tau* knockout recombined with the *K394R* mutant (*tau^K/O^*, *K394R*/*tau^K/O^*, *K394R*). **(b)** Quantification of crossing frequencies as probabilities of either (1) Partial Crossing or (2) Complete Crossing phenotypes being different from *w^1118^* control mushroom bodies. The frequency of Partial Crossing and Complete Crossing was higher in *tau^K/O^*/*tau^K/O^*mushroom bodies compared to *w^1118^* brains. Sample sizes: *w^1118^*= 35; *K394R/K394R* = 37; *tau^K/O^*/*tau^K/O^*= 41; tau^K/O^, *K394R*/ *tau^K/O^*, *K394R* = 41. Bars on graphs represent 95% confidence intervals (upper/lower ranges) for probabilities. Statistical significance indicated by different letters where “Partial Crossing” and “Complete Crossing” are treated independently from one another. Blue dashed boxes represent zoomed-in views of mushroom body midline. Scale bars: 50 μm, 10 μm.

## DISCUSSION

The microtubule cytoskeleton plays an essential role in shaping the structure of neurons, and its function is regulated by post-translational modifications like acetylation. Although multiple tubulin sites have been shown to be acetylated over the past several decades (L’Hernault and Rosenbaum 1985; Choudhary et al. 2009; Chu et al. 2011; Janke and Bulinski 2011; Janke and Magiera 2020), understanding the role acetylation plays in regulating neuronal microtubules is a constantly growing area of research. Previous work in our lab has supported the critical role that acetylation of α-tubulin K394 plays in regulating microtubules to shape axon terminal morphogenesis at the NMJ (Saunders et al. 2022). The results of our current study indicate that K394 acetylation is also important for the development of the mushroom body in the fly brain. Our results reveal that *K394R* homozygous mutant mushroom bodies show overextension of the β lobes at the brain midline. The *K394R* mutation does not overtly disrupt other mushroom body lobes, and our data suggest that its effect on β lobe growth is cell autonomous. Our findings support the idea that acetylation at K394 plays a critical role throughout *Drosophila* development, at both the NMJ during larval stages and in the central nervous system during metamorphosis.

In addition to analyzing the *K394R* mutant, we also attempted to disrupt K394 acetylation by manipulating HDAC6, which our previous findings indicate is the likely deacetylase that acts on K394 (Saunders et al. 2022). To determine whether and how HDAC6 acts during mushroom body development, we elevated HDAC6 levels and also used an *HDAC6* knockout allele to analyze how β lobe growth is affected. We found that overexpressing *HDAC6* alone increased the frequency of complete midline crossing, similar to *K394R*, and that the overexpression of *HDAC6* in the *K394R* homozygous mutants resulted in a complete crossing frequency that was equivalent to *HDAC6* overexpression or the *K394R* homozygous mutant alone. These data indicate that HDAC6 influences mushroom body development and are consistent with the model that the *K394R* mutant affects β lobe extension by disrupting K934 acetylation. While elevating HDAC6 perturbed β lobe development, the *HDAC6* knockout brains, in which K394 acetylation may be increased, had normal β lobes. This suggests that elevating K394 acetylation does not disrupt axon growth, although further testing of this idea depends on the identification of the K394 acetyltransferase (our previous studies indicate this is not αTAT, the acetyltransferase that acts on K40) (Saunders et al. 2022). In interpreting these HDAC6 experiments, it is important to keep in mind that HDAC6 is a multi-functional enzyme that targets multiple proteins as well as multiple α-tubulin sites, including the well-characterized K40 site (Valenzuela-Fernandez et al. 2008; Liu et al. 2015a). Thus, while our data are consistent with the model that HDAC6 exerts its effect on β lobe development by affecting K394 acetylation, it is possible that HDAC6 is also acting on additional proteins.

In the developing mushroom body, axons that compose the γ and α’/β’ lobes act as pioneers for α/β lobe development (Lee et al. 1999). In the *K394R* homozygous mutants, we found that extension of both the γ and β’ lobes was similar to controls. Although our data cannot eliminate the possibility that signaling from these other lobes to the β lobe is disrupted, our results suggest that *K394R* affects β lobe development in a cell-autonomous manner. It is possible that disrupting K394 acetylation in the *K394R* mutant causes β lobe overextension by impinging on axon guidance cues, such as Robo2/3, or through a signaling pathway, such as TGFβ (Ng 2008; Kang et al. 2019; Marmor-Kollet et al. 2019; Wu et al. 2021; Lin 2023) during early pupal stages in the *K394R* mutant.

While it is not clear specifically how K394 acetylation affects microtubules during mushroom body development, our experiments with *tau* provide some insightful information. Our data support a model in which mushroom body development is regulated by an interaction between the acetylation state of K394 and Tau. We found that elevating the levels of Tau caused an overextension of the β lobes and, in ∼20% of brains, perturbed the morphogenesis of the α and β lobes. Eliminating K394 acetylation via *K394R* increased the effects of *tau* overexpression, resulting in ∼90% of brains showing abnormal mushroom bodies—a striking 70% increase in the frequency of the phenotype. This finding was contrary to our initial expectation that we would observe a suppression of the *K394R* phenotype with elevated levels of Tau, similar to our finding that the overexpression of Futsch, which stabilizes microtubules, suppresses the *K394R* phenotype at the NMJ (Saunders et al. 2022). Our previous studies indicated that K394R decreases microtubule stability, and we had anticipated that Tau might act similarly to Futsch and stabilize microtubules in the *K394R* mutant. However, our results are consistent with recent reports suggesting that Tau may not (directly) stabilize microtubules (Qiang et al. 2018; Baas and Qiang 2019; Ori-McKenney and McKenney 2024). Instead, these new findings indicate that Tau affects microtubule structure to regulate the binding of other MAPs and that Tau may act to preserve labile domains, promoting dynamic states along axonal microtubules—in contrast to the typical microtubule-stabilizing function that Tau has been thought to have. In keeping with the idea that K394R and Tau may have opposite effects on microtubules (or may even negatively regulate each other), the β lobe axon extension defects observed in the homozygous *K394R* and *tau^K/O^*single mutants were eliminated in the *K394R* and *tau^K/O^*double-mutant. Altogether, our data support an interaction between K394 acetylation and Tau during axon morphogenesis in the fly brain.

## CONCLUSION

Microtubules play a central role in the morphology and function of neurons, and the acetylation of microtubules plays an important part in their regulation. Although it has been long known that tubulin is acetylated, only a single site, K40, has been extensively characterized. We have previously shown that acetylation of another conserved site, K394, is essential for the growth of axon terminals at the *Drosophila* larval NMJ, and our findings here show that it is also critical for the extension of axons that form the β lobes in the mushroom body of the adult brain. Our data suggest that there is an interaction between K394 acetylation and Tau that impacts mushroom body development. This study—combined with our previous studies—reveals that K394 acetylation is crucial at both the larval and pupal stages of fly development, and the resulting phenotypes are detectable into adult stages. The genetic tractability and high levels of gene conservation make fruit flies a valuable *in vivo* model to continue studying not just K394, but additional acetylation sites in order to uncover the effects of post-translational modifications on microtubule behavior and neuronal function. Furthermore, because disruptions in axon guidance are a cellular phenotype of neurodevelopmental disorders, our finding that *K394R* perturbs the growth of axons in the fly mushroom body suggests that this mutant may be a useful starting point to gain further insight into the misregulation of microtubules that contributes to neurological disease.

## DATA AVAILABILITY STATEMENT

The authors affirm that all data necessary for confirming the conclusions of the article are represented fully within the article and its figures.

## ACKNOWLEDGEMENTS

We thank the Bloomington *Drosophila* Stock Center and Dr. Sebastian Rumpf (Universität Münster) for fly strains and the Developmental Studies Hybridoma Bank for antibodies. We thank Monsserrat Pallan for technical assistance, Drs. Michael Perry (University of California, San Diego) and Kimberly Mulligan (California State University, Sacramento) for experimental advice, guidance, and feedback. We thank Angeli Hernandez (University of California, San Diego) for staff support, FlyBase, and members of the Wildonger laboratory for thoughtful input and scientific discussion.

